# Notes on the data quality of bibliographic records from the MEDLINE database

**DOI:** 10.1101/2022.09.30.510312

**Authors:** Robin Bramley, Stephen Howe, Haralambos Marmanis

**Affiliations:** Copyright Clearance Center Limited, London, United Kingdom; Copyright Clearance Center Inc., Danvers, Massachusetts, United States of America

## Abstract

The US National Library of Medicine has created and maintains the PubMed^®^database, a collection of over 33.8 million records that contain citations and abstracts from the biomedical and life sciences literature. That database is an important resource for researchers and information service providers alike. As part of our work related to the creation of an author graph for coronaviruses, we encountered several data quality issues with records from a curated subset of the PubMed database called MEDLINE. We provide a data quality assessment for records selected from the MEDLINE database and report on several issues ranging from parsing issues (e.g., character encodings and schema definition weaknesses) to low scores against several data quality metrics (e.g., identifier completeness, validity, and uniqueness).

## 1 Introduction

PubMed is an enormously valuable resource for the biomedical and health fields. The PubMed database is a voluminous collection of medical literature citations that is free, easily accessible, and has been a data source for many works in the information retrieval and life sciences communities. As machine learning becomes more prevalent in various branches of the life sciences, the number of works that rely on the PubMed database increases. Many papers that cited PubMed have appeared within the proceedings of The International Conference on Data and Text Mining in Biomedicine series e.g., DTMBIO ‘10 [1]. In ACM’s Digital Library[2], the year 2021 was a new high point at 235 for computing research articles that mentioned PubMed in the full-text collection, up from 1 in 1998 and 115 in 2010. Many information providers utilize the PubMed database, and there are a variety of machine learning models trained on PubMed[3]. It should be no surprise that, during the COVID-19 pandemic, the PubMed database has been crucial in providing timely and frictionless access to the scientific literature[4].

However, the PubMed database, which contains over 33.8 million records [5] collected over many decades, suffers from several data quality issues. These issues relate to, in part, character encodings, the absence of persistent identifiers, differences in human languages, and schema changes. These shortcomings should not be surprising since PubMed aggregates information produced by different publishers and XML providers, a fact that leads naturally to the presence of “multi-source problems” [6].

MEDLINE is a curated subset of PubMed, its records are indexed with a controlled vocabulary called MeSH [7] and include information regarding funding, genetic, chemical, and other metadata. Articles in MEDLINE predominantly come from a set of indexed journals and a reference data file of these journals is available separately [8]. MEDLINE was made available online, through PubMed, in 1997.

In this article, we will provide an account of our experience in working with the curated MEDLINE records and report on the data quality issues that we encountered. We will describe, at length, the problem of Author Name Disambiguation, which is widely acknowledged as a source of errors when processing bibliographic databases in general, due to the challenges of synonyms (e.g., “John Doe”, “John T Doe”, and “JT Doe” referring to the same individual) and homonyms (i.e., two different people who share the same name such as “John Smith”) [9]. Other problem areas that we will discuss include issues with character encodings, date related issues, the presence of persistent identifiers (and lack thereof), affiliation disambiguation, language related data issues, and schema data quality issues. Knowing how to address these challenges is valuable for practitioners who need to work with MEDLINE (or databases like MEDLINE) and process its records so that they can be used in their information systems.

### 1.1 PubMed data

The PubMed database is available as XML, based on a DTD (currently the 2019 version) [10]. The compressed files are made available via an FTP server (they are also accessible by HTTPS) and each one of them contains up to 30,000 citation records. Every year, in mid-December, the data are consolidated and an annual baseline is produced. This is followed by incremental daily update files that include deletions.

A PubMed XML file has a root element of `PubmedArticleSet` that contains 1, or more, `PubmedArticle` or `PubmedBookArticle` children. The DTD also permits 0 or 1 `DeleteCitation` elements, and these can be seen in the update files. The elements of the `PubmedArticle` are divided into the `MedlineCitation` and the optional `PubmedData` - we have colloquially referred to these as the “front” and “back” matter respectively.

The description of the XML elements [11], also outlines potential discrepancies caused by schema changes, or policy changes to the collected data. For example, records created before 2002 only contained author initials instead of full, first or middle, names; moreover, records between 1988 and 2013 only included the affiliation for the first author.

#### 1.1.1 Known DTD shortcomings

There are two known problems with the DTD that have not yet been addressed. The first known problem is that authors cannot be linked to their `CollectiveName`. Some publishers have tried to work around this by interspersing `CollectiveName` elements and `Author` elements. In a wheat genome sequencing consortium paper (PMID 30115783), one of the contributors was a member of 12 groups, so that person appears as an Author record 12 times. This multiplicity complicates the author name disambiguation, as it may be impossible to distinguish a duplicate author entry from a valid homonym.

The second problem is related to a shortcoming in the 2019 DTD. Specifically, the back matter `PubmedData` element may contain a `ReferenceList` with many `Reference` elements, but it doesn’t prevent the presence of many `ReferenceList` elements each with one `Reference`. Consequently, extraction must be able to handle both because both have been observed in the records. Furthermore, the `ReferenceList` definition permits deeply nested `ReferenceList` elements, as shown below:

~~~
<!ELEMENT ReferenceList (Title?, Reference*, ReferenceList*)>
~~~

#### 1.1.2 Escape characters

Escape sequence characters may appear within text fields such as the article title or abstract text. For example, if you wanted to represent a record in JSON, then you would have to beware of trailing backslashes and double quotes. Backslashes can also be problematic for the language used to parse the record. Furthermore, it may be necessary to remove other special characters such as new line characters (e.g., carriage return, line feed), tabs, and so on.

#### 1.1.3 Extended characters

PubMed encompasses articles published in many different languages, sometimes multiple languages. Consequently, fields such as the affiliation string, or parts of the author’s name, may contain extended characters. This is an important consideration for the disambiguation of author names.

### 1.2 Open Source libraries

Since PubMed has been a canonical source of biomedical citations, there are open source libraries to assist with parsing the records. Whilst none of these libraries were appropriate for our needs, they are included here for completeness.

For Python, pubmed_parser [12] is an active project, but only handles a constrained field list. The pymed [13] project, which is now archived, only parsed and cleansed a limited subset of the fields. It also seems that the design was intended to wrap the API.

For Java, there is pubmed-parser [14], which is based around the Java Architecture for XML Binding (JAXB). This project only had a short flurry of commits over 6 days in April 2021, consequently it is unclear whether this is actively maintained.

## 2 Materials and Methods

This work will identify challenges that can be faced when working with the MEDLINE data and categorize them along several dimensions of data quality [15].

### 2.1 Data acquisition

The PubMed baseline files were downloaded from their respective NLM FTP folders [16][17] and uploaded to separate folders on an S3 bucket.

### 2.2 Data processing

Figure 1 illustrates our data processing approach. The PubMed gzipped XML files were processed using Apache Spark 3.1.1 on Amazon EMR 6.3.1. The initial ingestion process extracted a few key properties, such as the PMID and DOI (from the `PubmedData` if present), before splitting the XML into two fragments representing the front matter (bibliographic metadata) and the back matter (references).

**Figure 1:**
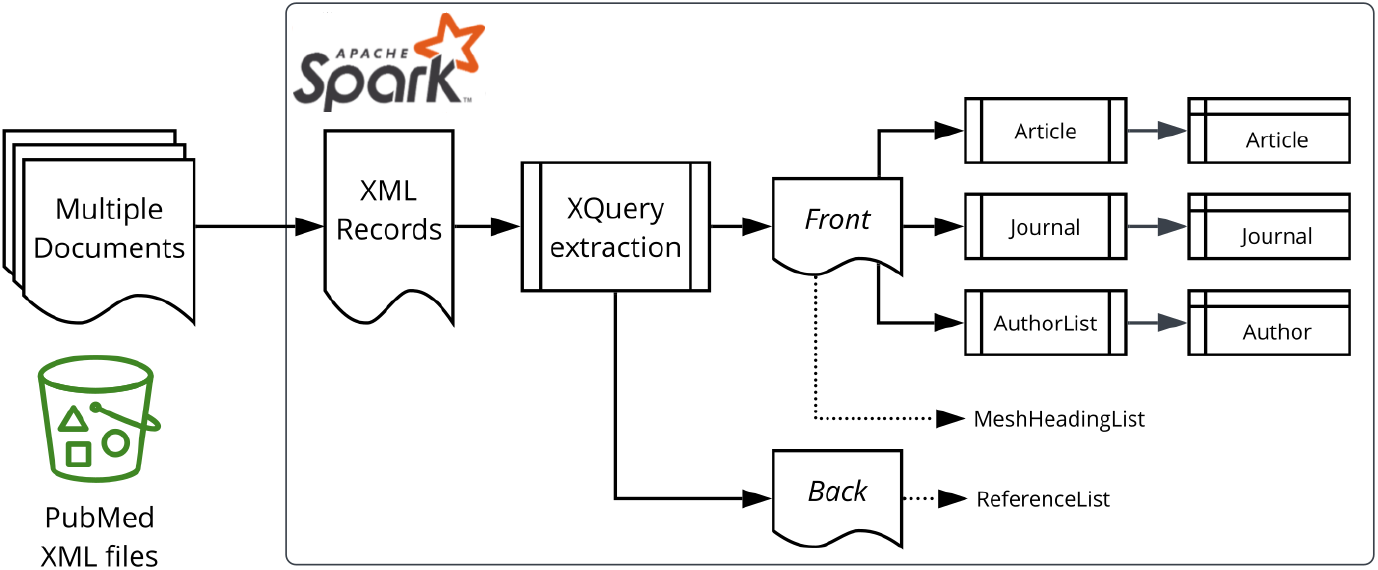
Data processing overview.

The baseline files were ingested first, then the update files were subsequently processed to apply updates, inserts and deletions. Record updates were applied by sorting the records by their PMID in conjunction with the `DateRevised` property; only the newest records were retained. Note that the PMID Version attribute is not suitable for this purpose as it is only used by Public Library of Science (PLOS) records [11].

Spark SQL [18] is designed for tabular data, with the key construct being the DataFrame. Whereas XML documents are represented using a hierarchical structure that allows for repeating elements (a one-to-many relationship). This leads to an inherent mismatch between the two data formats that requires data transformation.

There is a spark-xml module [19], but we discovered during our initial experiments that the PubMed XML was too complex for spark-xml, as it resulted in heavily nested DataFrames, and incorrect query results. Consequently, we solved the XML to DataFrame impedance mismatch by performing an XQuery [20] operation per target entity type (e.g. Article, Author, etc.) as shown on the right-hand side of Figure 1.

The spark-xml `XmlInputFormat` class was retained for loading the XML files into Spark, with the ingestion and extraction utilizing XQuery queries to extract properties, via the Saxon-HE [21] library as provided by the Elsevier Labs spark-xml-utils [22] module.

To ease maintenance of the complex XQuery queries, we adopted a pattern whereby the XQuery output produces a JSON document. This makes the target property for a particular XPath or XQuery expression transparent (Figure 2) and inserting new elements does not break downstream code because it does not rely on positional information. The last part of that transformation phase is to leverage the read method of the `SparkSession` object which parses the JSON documents to `DataFrame` records. Note that Figure 2 also represents the handling of escape characters using the XQuery `replace` function.

**Figure 2:**
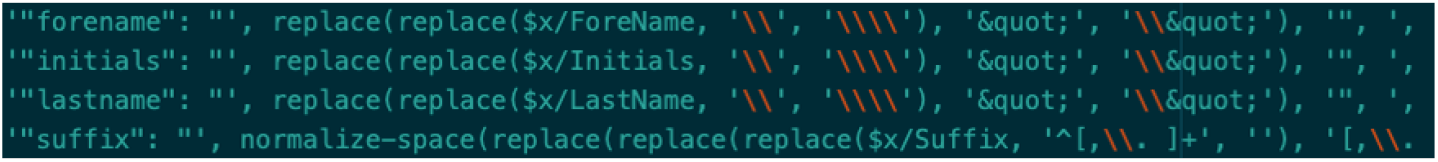
JSON representation within XQuery.

### 2.3 Data analysis

The resulting DataFrames were analyzed using Spark SQL in Apache Zeppelin [23]. For string fields, we consider the length in characters and in words (by splitting on spaces). Metrics were rounded to 3 decimal places (or less).

The plots were produced in R, with the box plots using log-scale for the y-axis.

### 2.4 Definitions

- 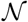 = number of records
- 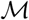 = number of records missing a value for the target property
- 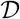 = distinct values of those present (excludes null / blank)
- 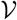 defined by count of records matching a regex for identifiers (Table 1)
- 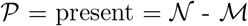
- *Completeness* metric = 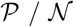
- *Validity* metric = 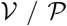
- *Uniqueness* metric = 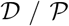

**Table 1:**
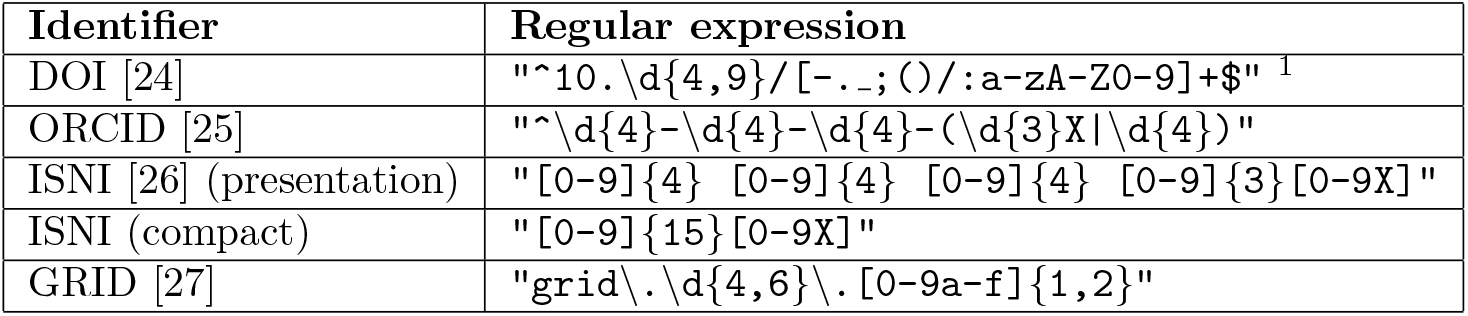
Regular expressions for identifier validation

### 2.5 Limitations of the study

The source dataset comprises the PubMed 2022 baseline plus daily update files to 1252 (30th March 2022).

It should be noted that our study includes only the `PubmedArticle` records, not the `PubmedBookArticle` records. The `PubmedArticle` records are only those from the MEDLINE subset (based on the `Status` attribute), and further excludes news articles, and those articles without a title; this gives a total of 28,986,590 article records. News articles were excluded from extraction because journalists, anecdotally those from the British Medical Journal, skew attempts to identify prolific authors through aggregation.

Other applied constraints are as follows:

- Only `Author` records with the `ValidYN` attribute of `Y` have been extracted, not `Investigator` records. For these 120,191,520 authors, only the first `Affiliation` element is considered.
- The `DataBank` element provides links to external datasets such as clinical trials. These identifiers were not investigated as part of the reported study.
- For alternative article identifiers, we did not extract the `ELocationID` element nor Publisher Item Identifiers (PII) from the `PubmedData`.
- For Journals, ISSNs were not analyzed.

#### 2.5.1 Approximation

Five number summary information is produced using Spark’s `DataFrameStatFunctions approxQuantiles` method with an error margin of 0.0001, an example is shown below:

~~~
articleDF.stat.approxQuantile(“doi_len”, Array(0.0,0.25,0.5,0.75,1.0), 0.0001)
~~~

However, the distinct counts do not leverage the Spark SQL `approx_count_distinct function`, rather the `dataframe.select(“column”).distinct.count` approach was used.

## 3 Results and discussion

In this section, we’ll present our results related to data quality for the entities and fields shown in Figure 3. The PubMed XML data model is article-centric, but we will work our way from left to right.

**Figure 3:**
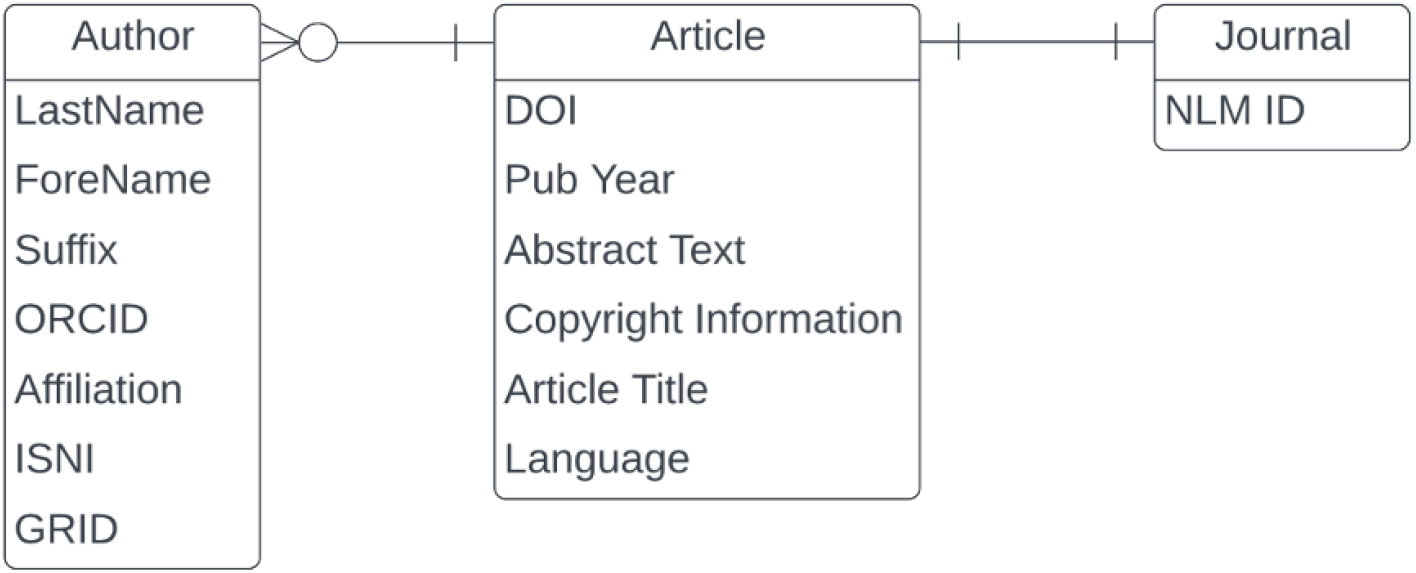
Entity Relationship Diagram for a subset of PubMed.

### 3.1 Data quality issues related to author names

One of the important considerations regarding author records is that PubMed has not always recorded all the authors of a paper. The number of authors was limited to 10 between the years 1984 and 1995, and to 25 between the years 1996 and 1999 [11].

The most common last names in MEDLINE are Romanized Chinese names (Table 2), which can be very challenging to disambiguate. Looking at the length characteristics (Figure 4), there are a few obvious problems, namely pollution of the author elements by incorrectly entered collective names (Table 3), and single character last names potentially caused by name transposition errors (Table 4).

**Figure 4:**
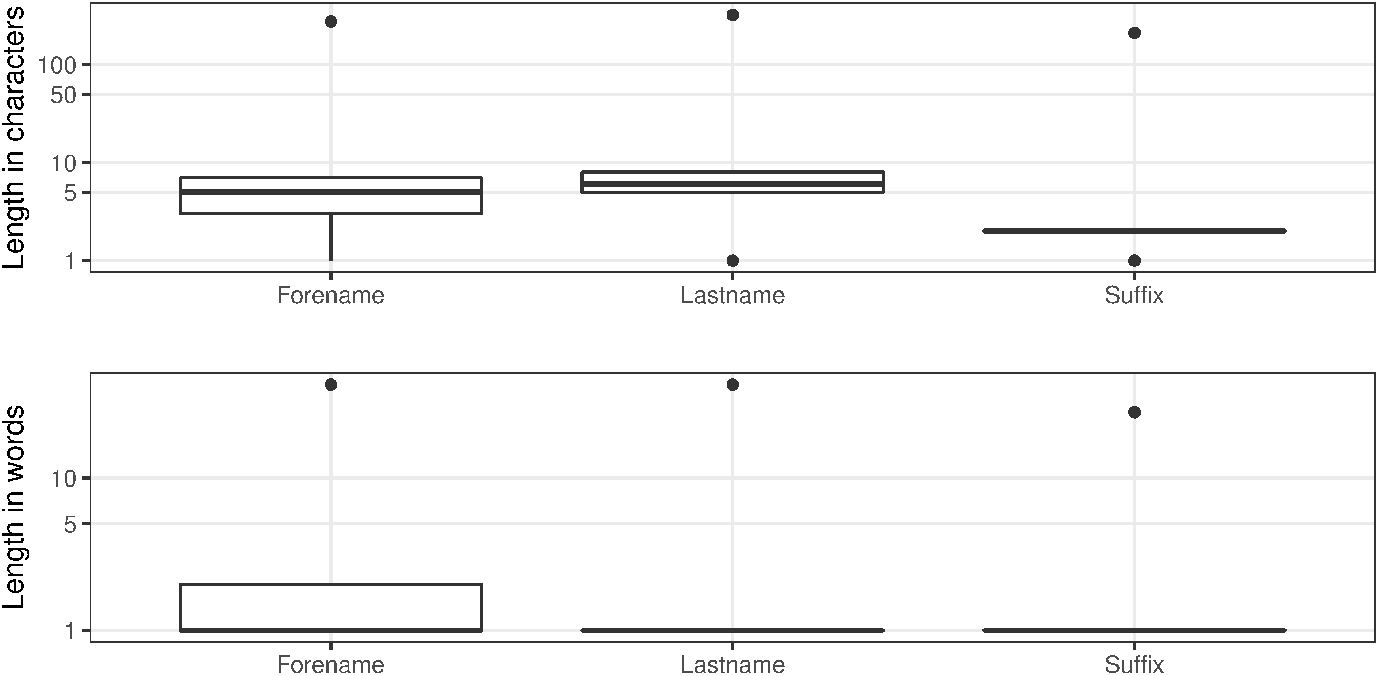
Author name character / word distributions.

**Table 2:** Top 10 LastName values.

| **LastName** | **Occurrences** |
| --- | --- |
| Wang | 1,086,073 |
| Li | 895,976 |
| Zhang | 878,544 |
| Chen | 722,753 |
| Liu | 703,743 |
| Lee | 547,636 |
| Kim | 523,687 |
| Yang | 433,439 |
| Wu | 360,532 |
| Huang | 309,375 |

**Table 3:** Ten longest LastName values.

| **LastName** | **Length** |
| --- | --- |
| Endocrinology Genetics And Metabolism Group Pediatric Branch Of Chinese Medical Association Neonatal Screening Group Specialist Committee For Prevention And Control Of Birth Defects Chinese Association Of Preventive Medicine Prevention And Control Committee Of Birth Defects Pediatric Branch Of Chinese Medical Association | 322 |
| The Group Of Minimally Invasive Spinal Surgery And Enhanced Recovery Professional Committee Of Orthopedic Surgery And Enhanced Recovery Association Of China Rehabilitation Technology Transformation And Promotion | 211 |
| Genetic Disease Society Guangdong Precision Medicine Application Association Prenatal Diagnosis Group Maternal And Child Health Care Society Guangdong Medical Association Expert Committee Of Prenatal Diagnosis | 209 |
| Arir Associazione Riabilitatori dell'Insufficienza Respiratoria Sip Società Italiana di Pneumologia Aifi Associazione Italiana Fisioterapisti And Sifir Società Italiana di Fisioterapia E Riabilitazione | 201 |
| This Paper Is A Co-Publication Between European Journal Of Preventive Cardiology European Heart Journal Acute Cardiovascular Care And European Journal Of Cardiovascular Nursing | 176 |
| Committee For Birth Defect Prevention And Control Chinese Association Of Preventive Medicine Genetic Testing And Precision Medicine Branch Chinese Association Of Birth Health | 174 |
| Consensus Group Of Experts On Application Of Metagenomic Next Generation Sequencing In The Pathogen Diagnosis In Clinical Moderate And Severe Infections | 152 |
| Expert Committee Of The Inter-Laboratory Quality Assessment Of Prenatal Screening And Diagnosis Clinical Test Center Of The National Health Commission | 150 |
| For The Antimalarial Therapeutic Efficacy Monitoring Group National Malaria Elimination Programme The Federal Ministry Of Health Abuja Nigeria | 142 |
| On Behalf Of The Association Of Rural Surgeons Of India-Lancet Commission On Global Surgery Consensus Committee Arsi-LCoGS Consensus Committee | 142 |

**Table 4:** Top 10 shortest LastName values.

| LastName | Occurrences |
| --- | --- |
| S | 756 |
| A | 704 |
| E | 636 |
| M | 592 |
| O | 563 |
| K | 497 |
| R | 453 |
| P | 363 |
| G | 306 |
| V | 279 |

The author forename field is 99.913% complete. Regarding the length, before 1945, the longest value in the forename field was 3 characters long, which reflects the policy to only hold author initials. The distributions, in Figure 4, clearly show that there are outliers. As shown in Table 5, these are primarily for working groups (a validity error), but the first row represents a different form of data preparation error where the affiliation has been concatenated with the forename.

**Table 5:** ForeName values over 100 characters.

| PMID | LastName value | ForeName value | Length |
| --- | --- | --- | --- |
| 34313229 | Choi | Moon Hyung Department Of Radiology Eunpyeong St Mary's Hospital College Of Medicine The Catholic University Of Korea Seoul Republic Of Korea Catholic Smart Imaging Center Eunpyeong St Mary's Hospital College Of Medicine The Catholic University Of Korea Seoul Republic Of Korea | 276 |
| 33145749 | En Representación Del Grupo de Trastornos de la Conducta Y Del Movimiento Durante El Sueño de la Sociedad Española de Sueño | En Representación Del Grupo de Trastornos de la Conducta Y Del Movimiento Durante El Sueño de la Sociedad Española de Sueño | 123 |
| 32329046 | En Representación Del Grupo de Estudio de Enfermedades Desmielinizantes de la Comunidad Autónoma de Madrid | En Representación Del Grupo de Estudio de Enfermedades Desmielinizantes de la Comunidad Autónoma de Madrid | 106 |
| 32433836 | Pharmakopsychiatrie | The Therapeutic Drug Monitoring Task Force Of The Arbeitsgemeinschaft Für Neuropsychopharmakologie Und | 102 |

Completeness does not apply to author suffixes since not everyone has a suffix to their name. In terms of uniqueness there are 823 distinct values across 483,541 entries. There are also consistency issues, examples of which can be observed in Table 6 (e.g., Jr, Junior, Júnior). Figure 4 shows the range of suffix lengths and clearly indicates that there is something wrong with at least some records. When we look at the longest values for author suffixes (Table 7) and the most common single character values (Table 8), it becomes clear that there are multiple data issues related to the author suffix field; the general theme of misplaced values, or value “pollution”, occurs across fields and is a major data quality weakness for the MEDLINE records.

**Table 6:** Top 15 suffixes.

| Suffix value | Occurrences |
| --- | --- |
| Jr | 374,510 |
| 3rd | 74,260 |
| 2nd | 20,364 |
| 4th | 5,828 |
| Sr | 4,075 |
| Junior | 535 |
| Júnior | 380 |
| Filho | 241 |
| PhD | 238 |
| 5th | 204 |
| Neto | 200 |
| III | 199 |
| Dr | 146 |
| 6th | 129 |
| MD | 99 |

**Table 7:** Ten longest suffixes.

| Suffix value | Length |
| --- | --- |
| Brian Buckley Caitlin Cornell Alyssa Fuller Eric Hojnowski Ryan LaFollette Yelena Livshits Todd Michaelis Claire Motyl Tarakad Ramachandran Devan Rahmachandrin Sofia Seckler Evaline Tso And Kate Zmijewski-Mekeem | 211 |
| European Society Of Clinical Microbiology And Infectious Diseases Escmid Vaccine Study Group Evasg | 98 |
| (Conceptualization; Review and editing; Read and approved final version of manuscript) | 86 |
| Faculty of Bioscience and Bioindustry, Tokushima University, Tokushima, Japan | 77 |
| BA, MBBS (Hons), FRANZCP, PhD, Dip Psychodynamic Psychotherapy, Cert ATP | 72 |
| on behalf of the Portuguese visual impairment study group (PORVIS-group) | 72 |
| (Writing original draft; Read and approved final version of manuscript) | 71 |
| RN, Cert Psych Nurs, BA (Hons), Dip Ed, B Ed, M Ed, PhD, FACMHN | 63 |
| DVM, PhD, Diplomate ABVP (Dairy Practice), SFHEA, NVS, MRCVS | 60 |
| B Phil (Hons), B Soc & Comm Stud (Community Development) | 60 |

**Table 8:** Top 10 shortest suffixes.

| Suffix value | Occurrences |
| --- | --- |
| * | 32 |
| S | 12 |
| K | 11 |
| W | 11 |
| J | 8 |
| F | 8 |
| † | 8 |
| A | 7 |
| P | 7 |
| M | 5 |

The PubMed DTD does not have a dedicated field for an email address. From 1996, NLM included “the first author’s electronic mail (e-mail) address at the end of <Affiliation>, if present in the journal. Furthermore, as of October 1, 2013, NLM no longer edits affiliation data to add e-mail address” [11]

A word of caution about relying on email addresses as a discriminator for author name disambiguation; the most common email address is which occurred 2023 times in the MEDLINE dataset of this study. Additionally, there are other non-specific email addresses such as journal editorial mailboxes.

Since 2010, the PubMed DTD has included an `Identifier` element, which has been used from 2013 [11]. However, it has less than 3% completeness (Table 9) and it is worth noting that there are occurrences where the same ORCID identifier has been incorrectly allocated to multiple authors within a paper.

**Table 9:** Author ORCID measures.

| Identifier | Completeness | Validity | Uniqueness |
| --- | --- | --- | --- |
| ORCID | 2.820% | 99.915% | 40.921% |

### 3.2 Data quality issues related to affiliation names

An author’s institutional affiliation is a very important information field, but the completeness is only around 42%. We have not derived a validity score, but there are quality problems within that set that are obvious from the length distributions (Figure 5). As previously mentioned, this field may contain values that aren’t written in English as well as non-ASCII characters.

**Figure 5:**
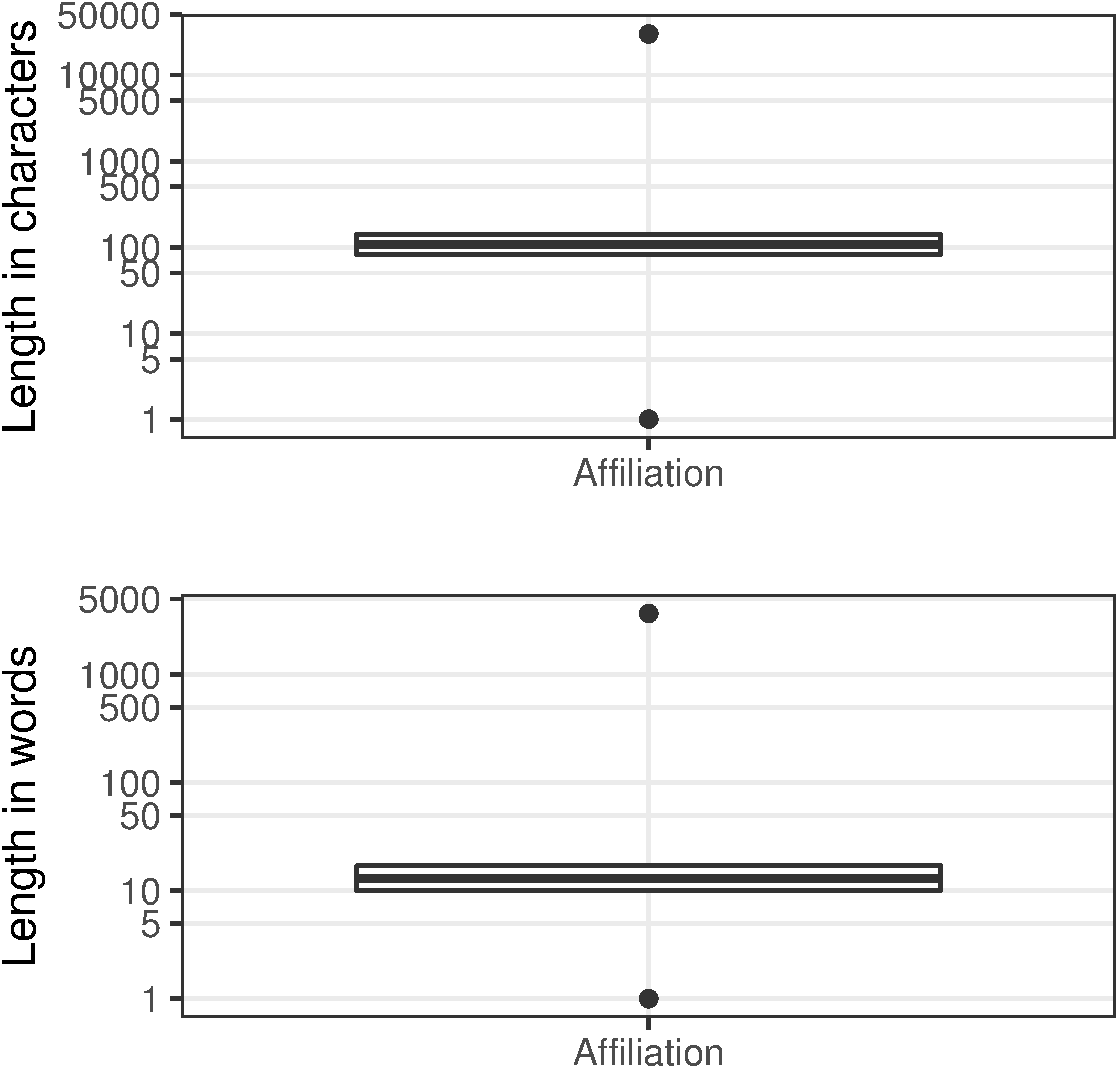
Affiliation character / word distributions.

In Figure 5, the outliers at the top of the range, which we have termed “narrative affiliations”, typically describe the affiliations for many, if not all, of the contributors to the paper (e.g., see Figure 6 where we show the entry from the article with PMID 32308221). These narrative affiliations may also be repeated for all the author entries within the author list. At the other end of the range, there are many incomplete, or indistinguishable entries (Table 10).

**Figure 6:**
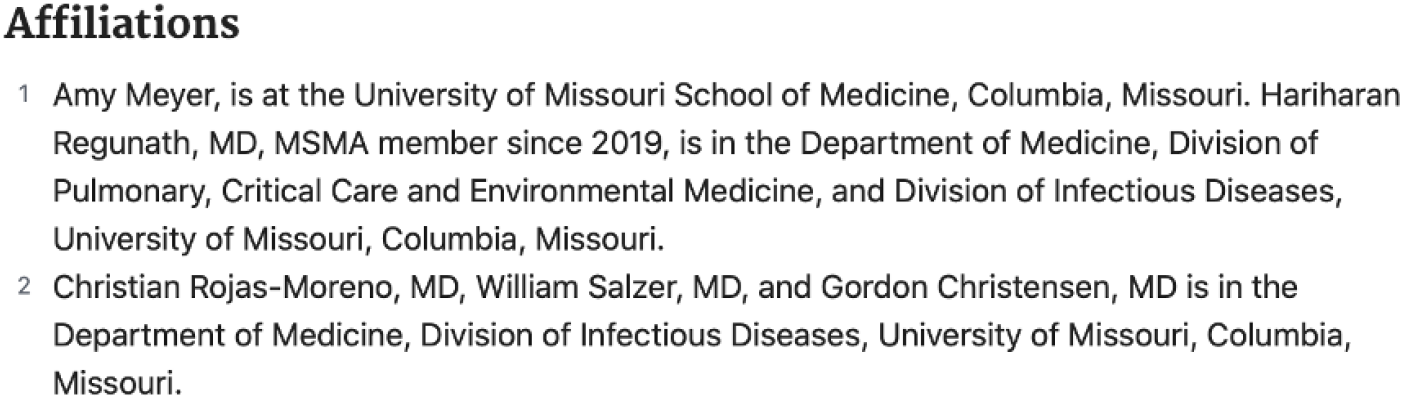
An example of narrative affiliations.

**Table 10:** Top 15 affiliations under 20 characters long.

| Affiliation string | Occurrences |
| --- | --- |
| . | 5,761 |
| ,. | 2,463 |
| London, UK. | 601 |
| Editor-in-Chief. | 468 |
| London. | 405 |
| Pathology. | 360 |
| GSK, Siena, Italy. | 342 |
| Duke University. | 341 |
| Harvard University. | 332 |
| McGill University. | 329 |
| Paris, France. | 323 |
| School of Medicine. | 303 |
| Yale University. | 301 |
| Editor. | 295 |
| Radiology. | 262 |

Our parsing has not included any special case exclusions. We note that pubmed_parser [12] excludes “For a full list of the authors’ affiliations please see the Acknowledgements section.” - though this exact string only occurs once within our selected dataset of over 51 million affiliation strings! It should also be noted that “as of October 1, 2013, NLM no longer performs quality control of the affiliation data” [11].

Whilst multiple affiliations were possible from the 2015 DTD [11], this is a good place to mention how some data providers concatenate multiple affiliations for an author in a single element. Here is an example for Yong-Beom Park (PMID 29465366):

> Division of Rheumatology, Department of Internal Medicine, Yonsei University College of Medicine, Seoul; and Institute for Immunology and Immunological Diseases, Yonsei University College of Medicine, Seoul, Republic of Korea.

Affiliation identifiers, such as ISNI and GRID, were possible from the 2015 DTD [11]. We’ve captured values for those too in Table 11.

**Table 11:** Key measures for Affiliations / Affiliation identifiers.

| Identifier | Completeness | Validity | Uniqueness |
| --- | --- | --- | --- |
| ISNI | 0.002% | 99.965% | 22.803% |
| GRID | 0.003% | 100.000% | 23.752% |
| Affiliation | 42.526% | N/A | 45.979% |

### 3.3 Data quality issues related to articles

#### 3.3.1 Article persistent identifiers

As can be seen in Table 12, the application of digital object identifiers (DOI), although not perfect, reaches a respectable score in terms of uniqueness but there are issues with validity of those identifiers and a significantly low score in terms of completeness; we’ll examine the impact that earlier publications have on DOI completeness.

**Table 12:** MEDLINE article identifiers.

| Identifier | Completeness | Validity | Uniqueness |
| --- | --- | --- | --- |
| DOI | 71.373% | 99.377% | 99.949% |

#### 3.3.2 Publication year

In the full PubMed database, there are over 100,000 records with a publication year earlier than 1900. In our selected data set from MEDLINE, there are only 3 that are clearly wrong (Table 13). In the first two examples, the publication year has the upper value from the journal pagination range. These erroneous publication years caused Parquet compatibility problems with Spark 3 (see issue SPARK-31404: https://issues.apache.org/jira/browse/SPARK-31404) when constructing a Date column, as they pre-date the introduction of the Gregorian calendar in 1582 and Spark implements a Proleptic Gregorian calendar as of version 3.

**Table 13:** Example of erroneous publication year values.

| PMID | Publication Year |
| --- | --- |
| 11662976 | 1132 |
| 11665278 | 1041 |
| 32422596 | 1 |

Figure 7 illustrates the volume of citation records with a valid DOI per publication year with 2022 in progress. Note that as of Q1 2022 there are not yet articles scheduled for publication in subsequent years.

**Figure 7:**
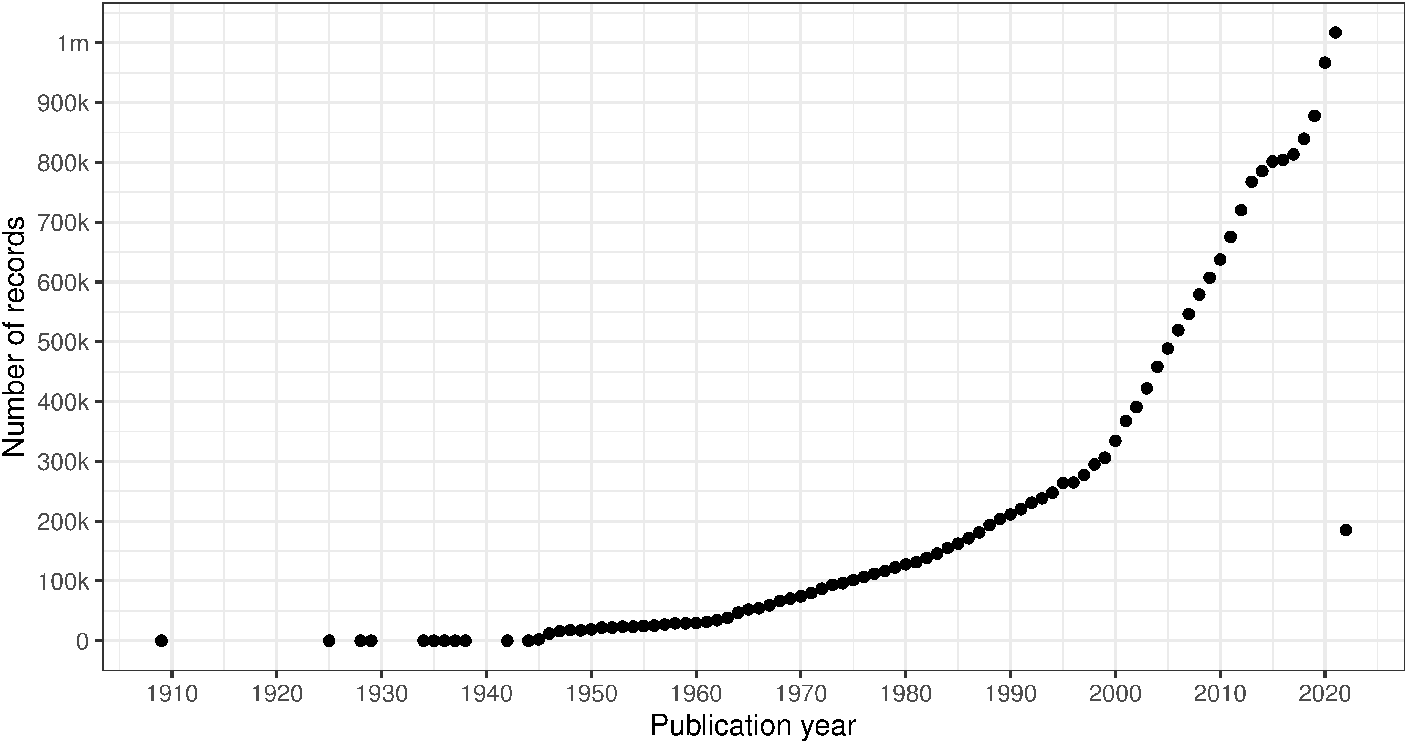
Count of citation records with a valid DOI per publication year (excluding erroneous years).

#### 3.3.3 Abstract

The abstract field was added to the PubMed record in 1975 [11]. The abstract text, which may be subject to copyright restrictions, is a prime candidate for text mining. Consequently, for the two-thirds of records with an abstract, it’s useful to understand their length distribution (Figure 8) and the erroneous values that they contain. Whilst the uniqueness is 99.942%, there is still a significant number (over 11 thousand abstracts) with non-unique abstract values. From the length information, we can infer that there are clearly meaningless abstract entries towards the lower end of these ranges, as seen in Table 14.

**Figure 8:**
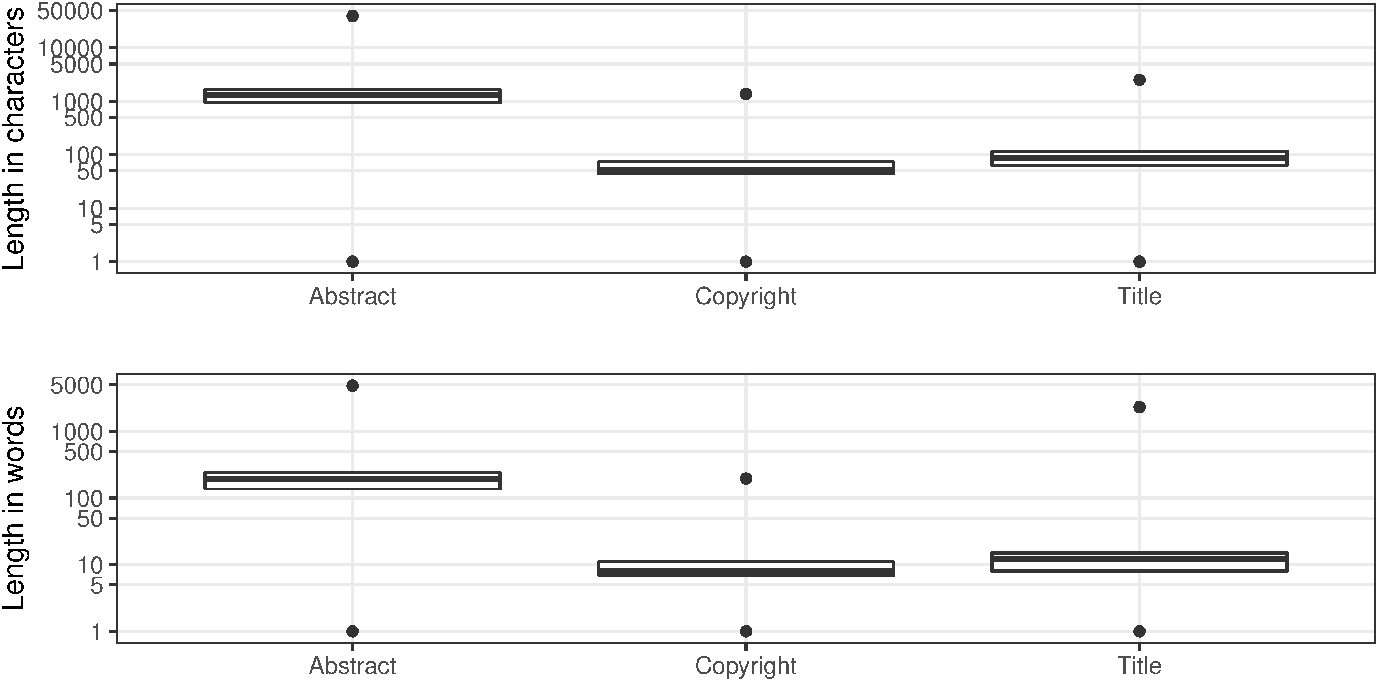
Article character / word distributions.

**Table 14:** Top 15 abstracts under 20 characters long.

| Abstract text | Occurrences |
| --- | --- |
| [Figure: see text]. | 579 |
| . | 182 |
| Not available. | 106 |
| N/A. | 51 |
| n/a. | 50 |
| no summary. | 48 |
| Null. | 41 |
| NA. | 29 |
| No Abstract. | 22 |
| <p/>. | 20 |
| Editorial. | 17 |
| EDITORIAL. | 16 |
|  | 13 |
| No abstract. | 13 |
| None. | 10 |

#### 3.3.4 Copyright

An important consideration when mining MEDLINE should be whether copyrighted material is being used. The NLM terms and conditions clearly state that they do not provide legal advice [28]. The copyright information field was introduced in 1999 [11], with a completeness measure of almost 22% of the records that have an abstract. From Table 15, it is evident that Elsevier is most consistent in supplying copyright statements although there is some lack of consistency regarding the actual values. Figure 8 shows the distributions of character length and word tokens, it should be clear that at the low end of the range there must be some invalid values (Table 16).

**Table 15:** Top 10 copyright statements.

| Copyright information | Occurrences |
| --- | --- |
| Copyright © 2020 Elsevier Inc. All rights reserved. | 40,773 |
| © 2021. The Author(s). | 39,577 |
| Copyright © 2020 Elsevier B.V. All rights reserved. | 39,221 |
| Copyright © 2020 Elsevier Ltd. All rights reserved. | 39,220 |
| Copyright © 2016 Elsevier Inc. All rights reserved. | 38,600 |
| Copyright © 2019 Elsevier Inc. All rights reserved. | 37,672 |
| Copyright © 2018 Elsevier Inc. All rights reserved. | 37,414 |
| Copyright © 2017 Elsevier Ltd. All rights reserved. | 36,833 |
| Copyright © 2017 Elsevier Inc. All rights reserved. | 36,817 |
| Copyright © 2018 Elsevier Ltd. All rights reserved. | 36,766 |

**Table 16:** Top 10 short copyright statements.

| Copyright information | Occurrences |
| --- | --- |
| © 2013. | 6,941 |
| excerpt | 4,996 |
| © The author(s). | 3,193 |
| © FASEB. | 1,444 |
| full text | 1,238 |
| ©2011 AACR. | 1,159 |
| ©2013 AACR. | 1,145 |
| ©2012 AACR. | 958 |
| Celsius. | 956 |
| © 2017 The Authors. | 925 |

#### 3.3.5 Title

MEDLINE has just over 7,500 records without an `ArticleTitle` element, leading to a completeness value of 99.974%. The uniqueness of the title field is approaching 98%. Like our observations for the abstracts, there are standard article titles that relate to the publication type towards the lower end of the character length and number of word token ranges (Figure 8; see also Table 17).

**Table 17:** Top 15 article titles under 20 characters long.

| Article title | Occurrences |
| --- | --- |
| [Not Available]. | 13,440 |
| Reply. | 1,972 |
| Invited commentary. | 1,896 |
| Editorial comment. | 1,676 |
| Editorial. | 1,465 |
| Response. | 1,312 |
| Discussion. | 1,052 |
| Editorial Comment. | 1,051 |
| Preface. | 974 |
| The authors reply. | 768 |
| In reply. | 714 |
| Introduction. | 585 |
| In Reply. | 519 |
| Authors' response. | 469 |
| Foreword. | 428 |

#### 3.3.6 Language

Another important consideration for text mining is the language, or languages, that the article is published in. It should be noted that PubMed includes translated titles, in square brackets, where appropriate. The language element contains language codes from the US Library of Congress MARC [29] schema, such as “chi” for Chinese. The language code table [30] includes *“und*” for undetermined and *“mul*” for multiple languages. However, language codes can also be concatenated together; for example, *“fregerita*” means the article was published in French, German, and Italian.

The language field is complete for the entirety of the MEDLINE records, but if we treat a solitary value of *“und”* or *“mul”* (238,470 and 1,399 occurrences, respectively) as invalid then the validity of this field is 99.55%. This excludes cases where they are present with other values too. From a recency perspective, *“und*” last occurred in 2002, and that is the only occurrence since 1985; *“mul”*occurred once in both 2016 and 2015, but before that it was last seen in 2011.

The maximum number of languages specified for a record is 6, but the 75th percentile is 1. Considering the values individually by splitting the strings and exploding the resulting array, allows us to produce the top 10 languages (Table 18). Note that almost 84% of records within the MEDLINE sample are published in English. The next most common language, German, only accounts for about 3% of articles.

**Table 18:** Top 10 languages.

| Language code | Occurrences |
| --- | --- |
| eng | 24,290,379 |
| ger | 861,109 |
| fre | 744,111 |
| rus | 697,806 |
| jpn | 429,283 |
| spa | 364,920 |
| chi | 329,153 |
| ita | 305,526 |
| und | 239,588 |
| pol | 172,956 |

### 3.4 Data quality issues related to journals

The key identifier provided in MEDLINE for a journal is the US National Library of Medicine (NLM) identity. When compared to the J_MEDLINE reference data set of MEDLINE indexed journals [8], the NLM identifiers have a referential integrity [15] measurement of 99.989%. There were 146 NLM identifiers that were not included within the J_MEDLINE dataset, affecting 3,045 articles. When considering a graph representation of the dataset, this would result in dangling edges that may not be permitted by some graph storage engines, such as Neo4j.

### 3.5 Data quality issues related to time evolution

In this section we consider the change over time for some of the key identifiers. Are there any obvious trends in whether identifiers are becoming more pervasive or prevalent in newer citation records? Here are some general observations: DOIs are almost ubiquitous for new articles (Figure 9), ORCIDs have been on the rise to just under 17% of authors per year (Figure 10), but GRID and ISNI usage peaked in 2017, having first appeared in 2015 (Figure 11). That leaves us with the tedious task of disambiguating the affiliation of the authors in the records. As can be seen in Figure 12, the vast majority of recent records contain an affiliation string for all authors; this is due to a policy change in 2014 to collect affiliations for all contributors [11].

**Figure 9:**
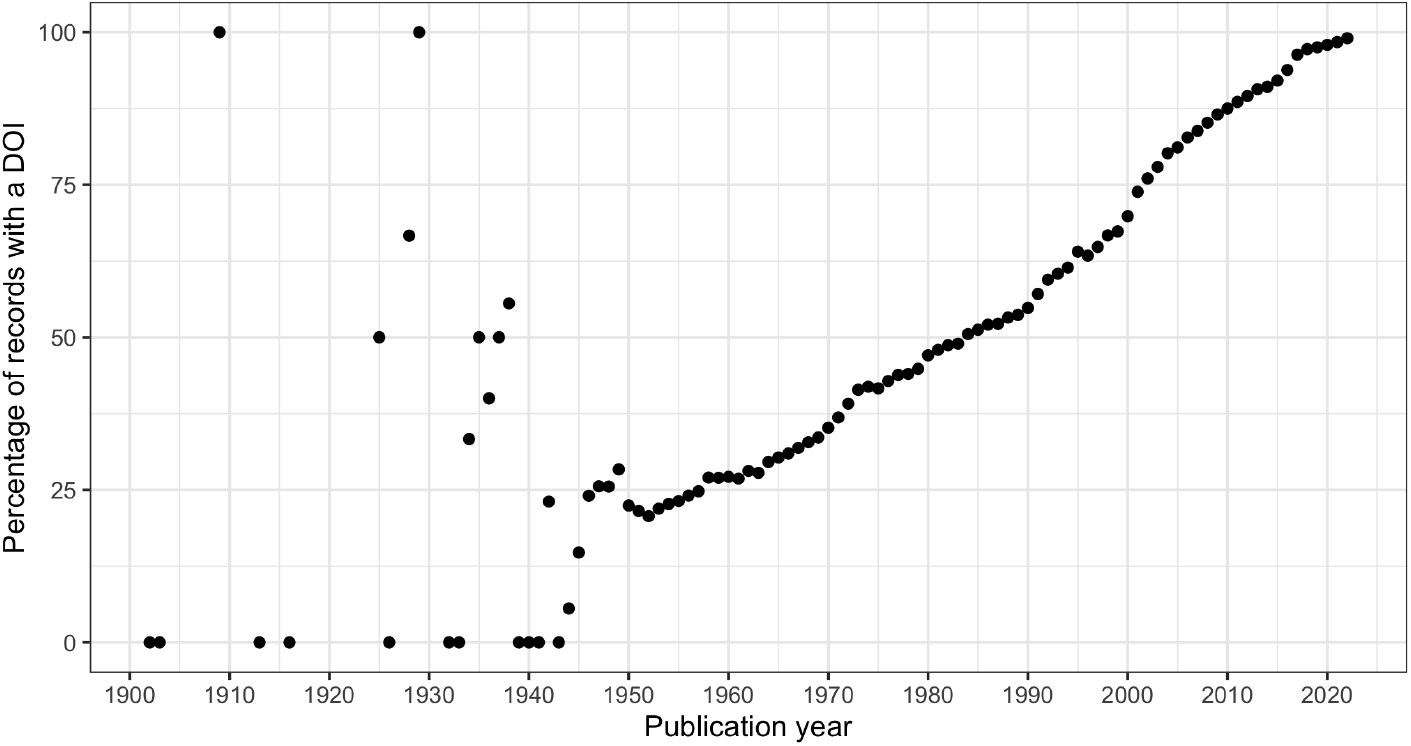
DOI percentage of articles per publication year.

**Figure 10:**
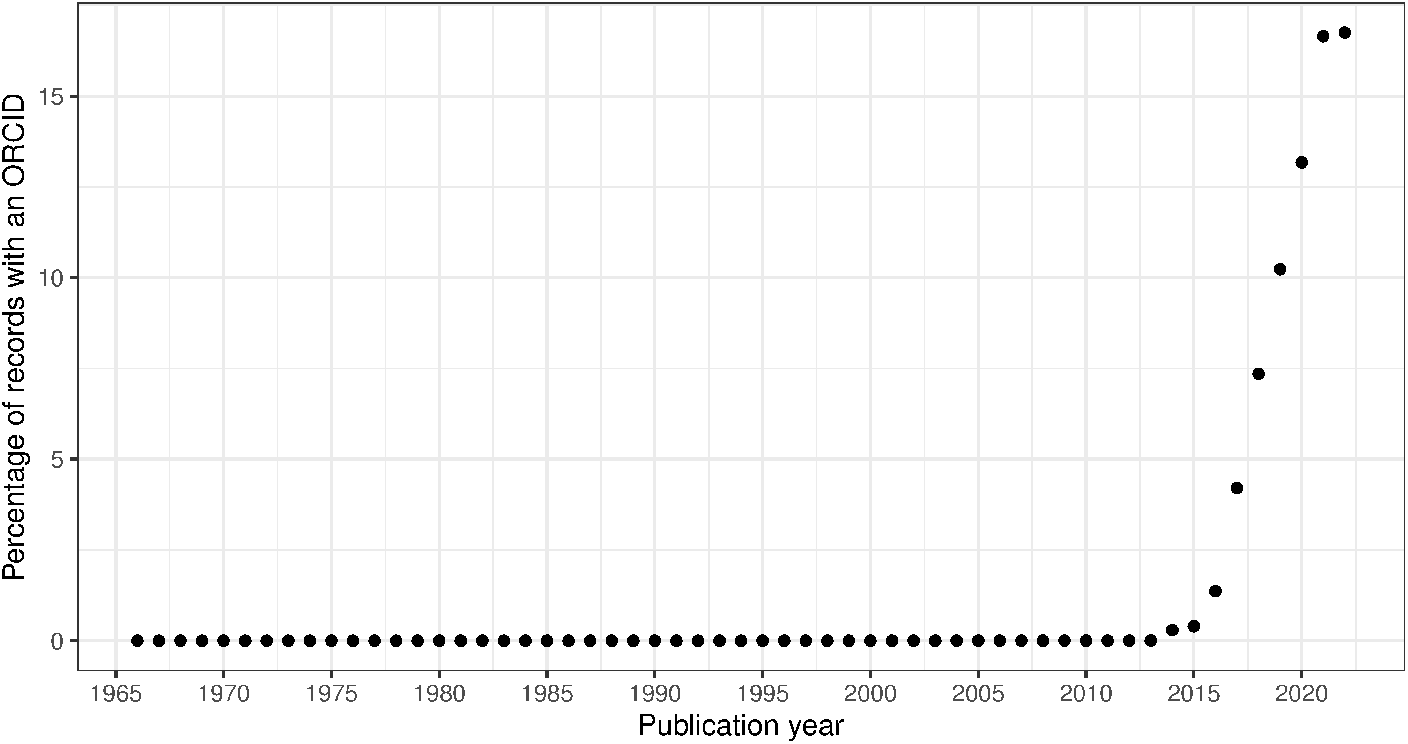
ORCID percentage of authors per publication year.

**Figure 11:**
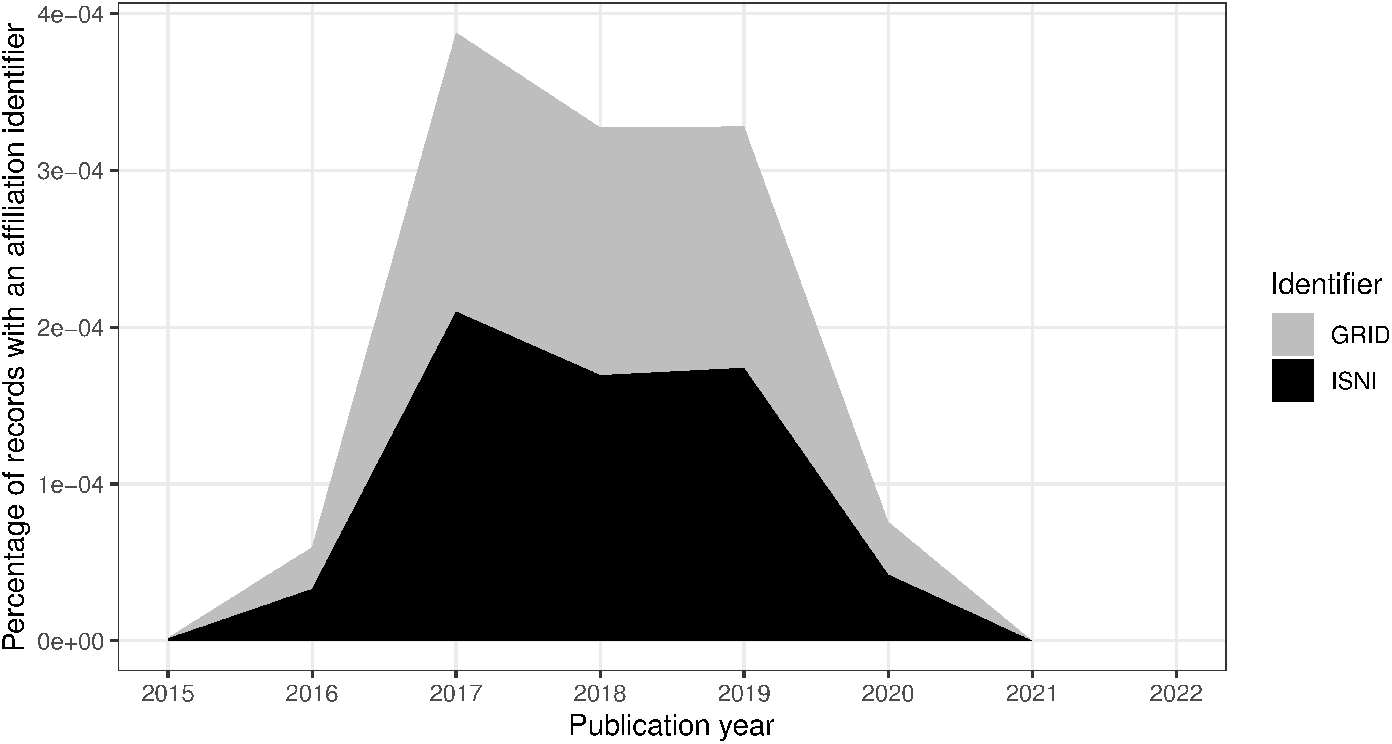
ISNI & GRID percentage of authors per publication year.

**Figure 12:**
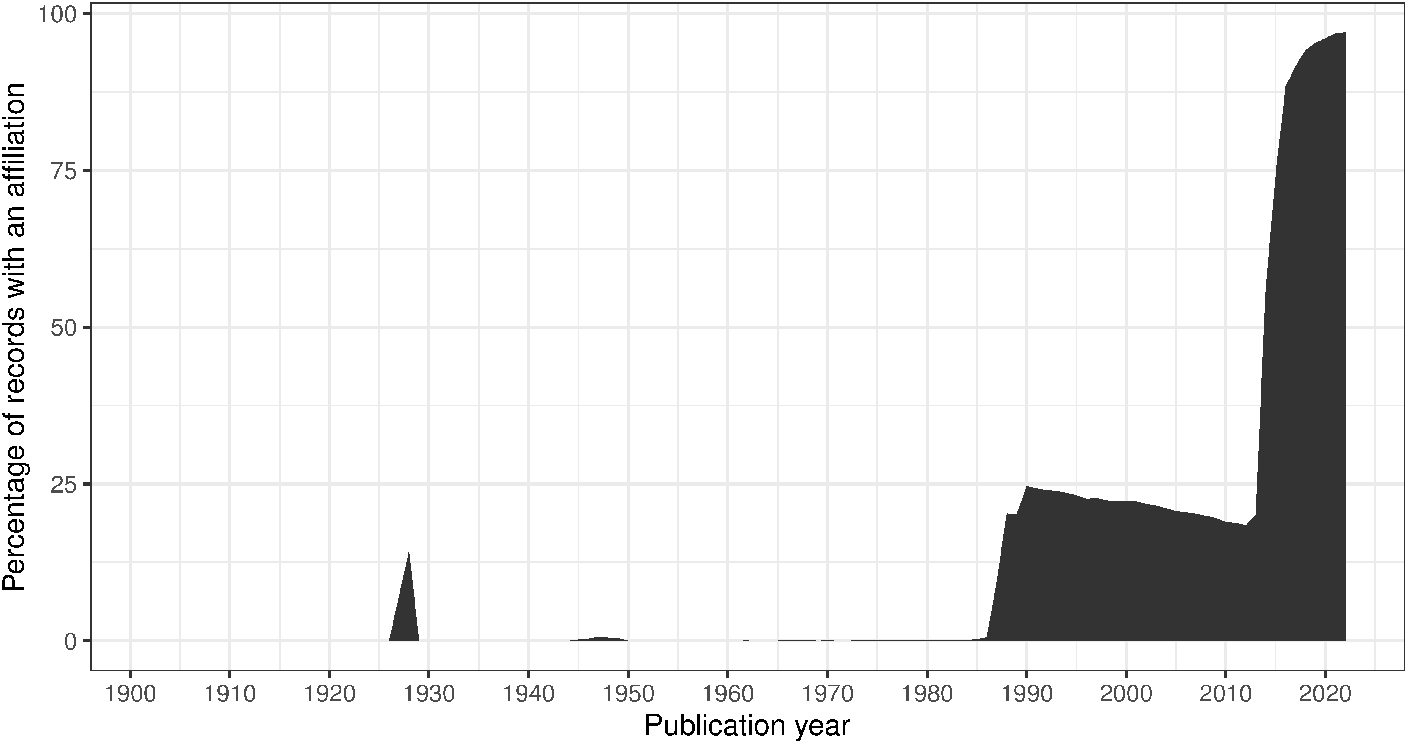
Percentage of authors per publication year with an affiliation string.

## 4 Conclusions

PubMed is an enormously valuable resource for the biomedical sciences and healthcare, yet, those attempting to identify authors and affiliations, or otherwise use the records from that database, need to be aware of the quality issues within the dataset. This article has highlighted some of those data quality concerns.

The data are subject to many human errors, such as typographical errors, and system related errors such as inconsistent representations of author names (leading to the synonym problem) and affiliations. There is a lack of author identifiers (contributing to the homonym problem) and a significant lack of affiliation identifiers. Being an aggregated source, the PubMed database suffers from multi-source problems such as inconsistent representations from the upstream XML providers that result in a high degree of lexicographic entropy.

In summary, our work supports the following conclusions:

- Given the incompleteness and uniqueness of identifying fields, the disambiguation of author names remains a significant problem for PubMed, particularly for records dating before 2014.
- PubMed has excellent integrity for NLM-internal identifiers (e.g., MeSH), though there is the noted exception around the J_MEDLINE dataset. Beyond the NLM database, the majority of articles are labelled with a DOI, and the DTD provides support for identifiers for authors, institutions, both of which are far from complete. The DTD also caters for grant information, and auxiliary data through the DataBank elements, though these were beyond the scope of our work.
- Overall, there is an improvement in the use of identifiers; in particular, records created since 2015 exhibit an increase in external identifiers. However, the data quality for institutional identifiers is poor and their use has been diminishing over time.

Unless the data quality issues are addressed retroactively, they will weaken (if not entirely distort) any subsequent data analysis. Perhaps, an intervention in current publishing systems, to prevent the data sources of PubMed from manifesting the data quality issues mentioned herein, is the best one can hope for the future. Much like the application of machine learning has been applied within the NLM for indexing (e.g., with the MTI tooling [31]), the NLM could enhance their process with systems that possess a learning architecture to improve and accelerate the curation of the PubMed records. It is also possible that another information provider will provide an open data repository containing cleansed PubMed data, although a proprietary offering is more likely.

Another possibility for better use of the PubMed treasure trove is the creation of an open source library for cleansing the data, or at least properly identify the data quality issues, and optimize the amount of information that one can obtain from processing the PubMed records. Once this is accomplished with one programming language the open source community can augment the library and expand its adoption in other programming languages, for example by porting the library.

Lastly, the community would benefit from the availability of open source libraries that can accurately perform author name disambiguation, or a substantial set of “gold data” that can be used for training and validation; that dataset, however, should be orders of magnitude larger than the ones that are currently available (e.g., the ‘amorgani/AND’ dataset [32] [33]).

## Footnotes

1 Adapted from https://www.crossref.org/blog/dois-and-matching-regular-expressions/

## References

[1] DTMBIO ‘10: Proceedings of the ACM fourth international workshop on Data and text mining in biomedical informatics. URL: https://dl.acm.org/doi/proceedings/10.1145/1871871.

[2] ACM Digital Library. URL: https://dl.acm.org.

[3] Lee J et al. “BioBERT: a pre-trained biomedical language representation model for biomedical text mining.” In: Bioinformatics 36.4 (2020), pp. 1234–1240. DOI: 10.1093/bioinformatics/btz682.

[4] LitCovid, NLM. URL: https://www.ncbi.nlm.nih.gov/research/coronavirus/.

[5] About PubMed, NLM. URL: https://pubmed.ncbi.nlm.nih.gov/about/.

[6] Rahm and Do. “Data Cleaning: Problems and Current Approaches”. In: Bulletin of the IEEE Computer Society Technical Committee on Data Engineering (2000).

[7] MeSH: Medical Subject Headings. URL: https://www.nlm.nih.gov/mesh/meshhome.html.

[8] Dataset of MEDLINE indexed journals. URL: https://ftp.ncbi.nlm.nih.gov/pubmed/J_Medline.txt.

[9] Sanyal DK, Bhowmick PK, and Das PP. “A review of author name disambiguation techniques for the PubMed bibliographic database”. In: Journal of Information Science 47.2 (2021), pp. 227–254. DOI: 10.1177/0165551519888605.

[10] PubMed 2019 DTD. URL: http://dtd.nlm.nih.gov/ncbi/pubmed/out/pubmed_190101.dtd.

[11] MEDLINE PubMed XML Element Descriptions and their Attributes. URL: https://www.nlm.nih.gov/bsd/licensee/elements_descriptions.html.

[12] Titipat Achakulvisut, Daniel E. Acuna, and Konrad Kording. “Pubmed Parser: A Python Parser for PubMed Open-Access XML Subset and MEDLINE XML Dataset XML Dataset”. In: Journal of Open Source Software 5.46 (2020), p. 1979. DOI: 10.21105/joss.01979. URL: https://doi.org/10.21105/joss.01979.

[13] pymed. URL: https://github.com/gijswobben/pymed.

[14] pubmed-parser. URL: https://github.com/thecloudcircle/pubmed-parser.

[15] DAMA International. DAMA - DMBOK Data Management Body of Knowledge. 2nd Edition. New Jersey, USA.: Technics Publications, 2017.

[16] PubMed baseline download files. URL: https://ftp.ncbi.nlm.nih.gov/pubmed/baseline/.

[17] PubMed daily update files. URL: https://ftp.ncbi.nlm.nih.gov/pubmed/updatefiles/.

[18] Michael Armbrust et al. “Spark SQL: Relational Data Processing in Spark”. In: Proceedings of the 2015 ACM SIGMOD International Conference on Management of Data. SIGMOD ‘15. Melbourne, Victoria, Australia: Association for Computing Machinery, 2015, pp. 1383–1394. ISBN: 9781450327589. DOI: 10.1145/2723372.2742797. URL: https://doi.org/10.1145/2723372.2742797.

[19] The spark-xml project. URL: https://github.com/databricks/spark-xml.

[20] XQuery 3.0 standard. URL: https://www.w3.org/TR/2014/REC-xquery-30-20140408/.

[21] Saxon-HE library, Saxonica. URL: http://www.saxonica.com/.

[22] The spark-xml-utils project. URL: https://github.com/elsevierlabs-os/spark-xml-utils.

[23] Apache Zeppelin. URL: https://zeppelin.apache.org.

[24] Digital Object Identifier (DOI). URL: https://www.doi.org.

[25] Open Researcher and Contributor ID (ORCID). URL: https://orcid.org.

[26] ISO 27729, International Standard Name Identifier (ISNI). URL: https://isni.org.

[27] Global Research Identifier Database (GRID), Digital Science. URL: https://www.grid.ac.

[28] NLM terms and conditions. URL: https://www.nlm.nih.gov/databases/download/terms_and_conditions.html.

[29] MARC definition, US Library of Congress. URL: https://www.loc.gov/marc/faq.html#definition.

[30] MEDLINE/PubMed Language Table. URL: https://www.nlm.nih.gov/bsd/language_table.html.

[31] NLM Medical Text Indexer (MTI). URL: https://lhncbc.nlm.nih.gov/ii/tools/MTI.html.

[32] Vishnyakova D, Rodriguez-Esteban R, and Rinaldi F. “A new approach and gold standard toward author disambiguation in MEDLINE.” In: J Am Med Inform Assn 26 (2019), pp. 1037–1045. DOI: 10.1093/jamia/ocz028.

[33] Vishnyakova D et al., AND - Author Name Disambiguation corpus. URL: https://github.com/amorgani/AND/.

